# Distributed brain co-processor for tracking electrophysiology and behavior during electrical brain stimulation

**DOI:** 10.1101/2021.03.08.434476

**Authors:** Vladimir Sladky, Petr Nejedly, Filip Mivalt, Benjamin H. Brinkmann, Inyong Kim, Erik K. St. Louis, Nicholas M. Gregg, Brian N. Lundstrom, Chelsea M. Crowe, Tal Pal Attia, Daniel Crepeau, Irena Balzekas, Victoria Marks, Lydia P. Wheeler, Jan Cimbalnik, Mark Cook, Radek Janca, Beverly K. Sturges, Kent Leyde, Kai J. Miller, Jamie J. Van Gompel, Timothy Denison, Gregory A. Worrell, Vaclav Kremen

**Affiliations:** Department of Neurology, Bioelectronics Neurophysiology and Engineering Laboratory, Mayo Clinic, Rochester, MN, USA; International Clinical Research Center, St. Anne’s University Hospital, Brno, Czech Republic; Department of Physiology and Biomedical Engineering, Mayo Clinic, Rochester, MN, USA; Center for Sleep Medicine, Departments of Neurology and Medicine, Divisions of Sleep Neurology & Pulmonary and Critical Care Medicine, Mayo Clinic, Rochester, MN; Department of Veterinary Clinical Sciences, University of California, Davis, CA, USA; Department of Neurology, Royal Melbourne Hospital, Melbourne, Australia; Cadence Neuroscience, Seattle, WA; Medtronic PLC, Minneapolis, MN; Department of Bioengineering, Oxford University, Oxford, UK; Department of Neurologic Surgery, Mayo Clinic, Rochester, MN, USA; Faculty of Electrical Engineering, Czech Technical University in Prague, Prague, Czech Republic; Czech Institute of Informatics, Robotics, and Cybernetics, Czech Technical University in Prague, Prague, Czech Republic; Faculty of Biomedical Engineering, Czech Technical University in Prague, Kladno, Czech Republic; Faculty of Electrical Engineering and Communication, Brno University of Technology, Brno, Czech Republic; The Czech Academy of Sciences, Institute of Scientific Instruments, Brno, Czech Republic; Biomedical Engineering and Physiology Graduate Program, Mayo Clinic Graduate School of Biomedical Sciences; Mayo Clinic School of Medicine and the Mayo Clinic Medical Scientist Training Program; Second Faculty of Medicine, Motol University Hospital, Charles University, Prague, Czech Republic

**Keywords:** epilepsy, seizures, electrophysiology, machine-learning

## Abstract

Early implantable epilepsy therapy devices provided open-loop electrical stimulation without brain sensing, computing, or an interface for synchronized behavioral inputs from patients. Recent epilepsy stimulation devices provide brain sensing but have not yet developed analytics for accurately tracking and quantifying behavior and seizures. Here we describe a distributed brain co-processor providing an intuitive bi-directional interface between patient, implanted neural stimulation and sensing device, and local and distributed computing resources. Automated analysis of continuous streaming electrophysiology is synchronized with patient reports using a hand-held device and integrated with distributed cloud computing resources for quantifying seizures, interictal epileptiform spikes, and patient symptoms during therapeutic electrical brain stimulation. The classification algorithms for interictal epileptiform spikes and seizures were developed and parameterized using long-term ambulatory data from 9 humans and 8 canines with epilepsy, and then implemented prospectively in out-of-sample testing in 2 pet canines and 4 humans with drug resistant epilepsy living in their natural environments.

## Introduction

Epilepsy affects nearly 1.0% of the world population and is associated with a high disease burden.^1,2^ Approximately 1/3 of people with epilepsy continue to have seizures despite antiseizure medications.^3^ Electrical brain stimulation has emerged as a reversible and effective palliative therapy for drug resistant epilepsy, but therapy optimization is slow and long-term seizure freedom rare.^4,5^ Despite the addition of brain sensing current electrical stimulation devices lack accurate seizure diaries.^6–8^ Currently physician rely on patient seizure diaries that are known to unreliable^9,10^ coupled with incomplete electrographic data.^6,8^ The challenge of patient management without accurate seizure counts has remained a persistent technology gap impeding epilepsy management.

Here we describe a distributed brain co-processor for integration of implantable brain sensing and stimulation devices with off-the-body commercial electronics for clinical and neuroscience research applications.^11–14^ The integration of implantable devices with commercial electronics via bi-directional wireless connectivity allows algorithm complexity to scale with advances in consumer computer hardware. Brain implants providing sensing and bi-directional wireless connectivity enable continuous electrophysiology data streaming, and when coupled with off-the-body computing resources overcome the computational and data storage limitations of current implantable electrical brain stimulation (EBS) devices. Until recently, there were several obstacles to consolidating the technology layers required for EBS, streaming continuous brain electrophysiology, and synchronized behavior reports. Here we utilize the investigational Medtronic Summit RC+S^™^ (RC+S^™^), a rechargeable sensing and stimulation implantable device with a bi-directional application programming interface, to demonstrate these capabilities in canines and humans living with epilepsy.^11–13,15^ The system enables continuous streaming of intracranial electroencephalography (iEEG) to a hand-held tablet or smartphone for real-time analysis and tracking of interictal epileptiform spikes (IES), seizures, and correlation with synchronized patient reports (Fig. 1). The electrophysiology classifiers (seizure and IES) were validated, tested, and then prospectively deployed for out-of-sample testing in pet canines and humans living in their natural environments with epilepsy.

**Fig. 1.**
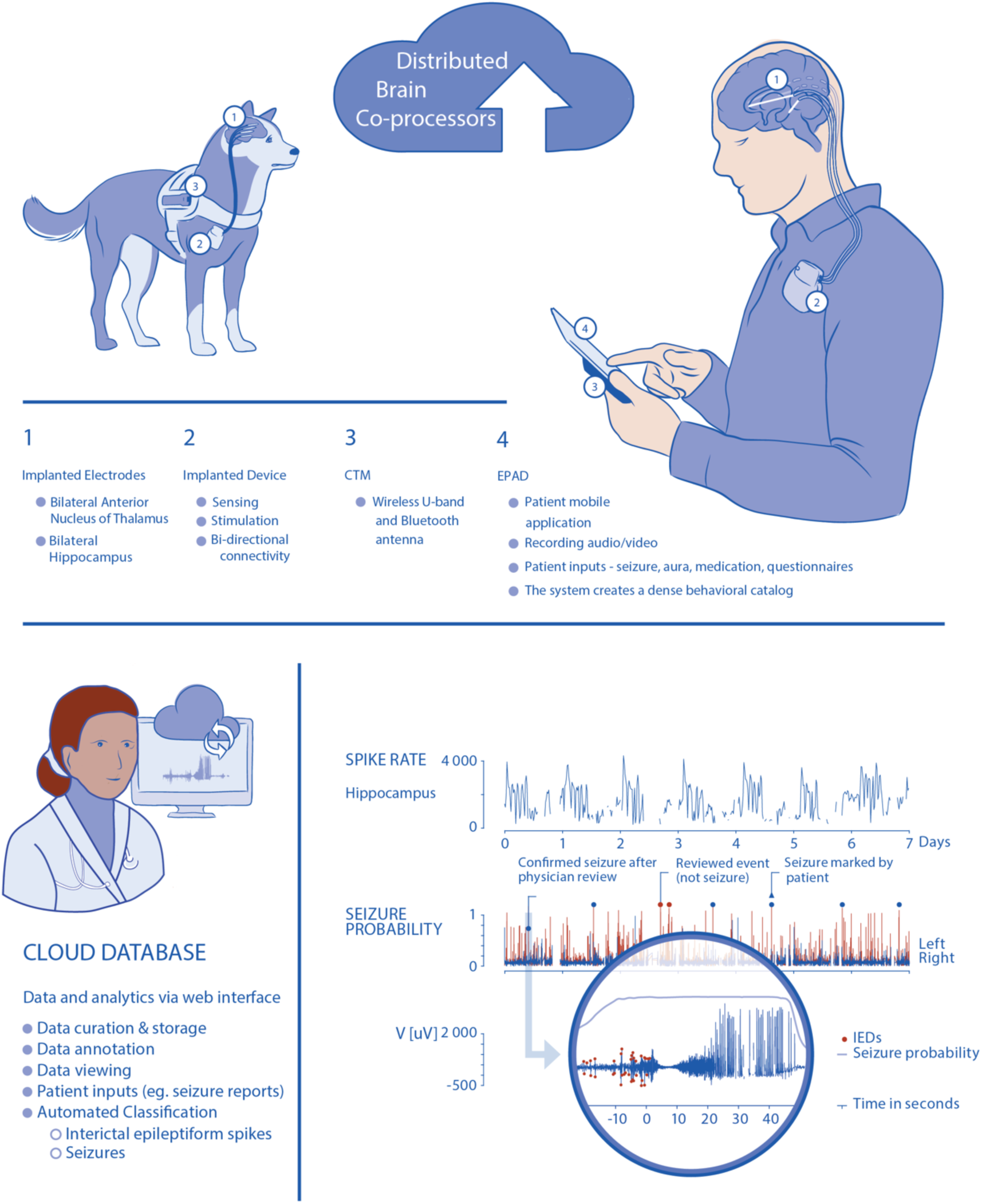
Distributed Brain Co-processor. Integrating implanted sensing and stimulation devices with off-the-body and cloud computing resources. The system was developed and prospectively tested in canines and humans with drug resistant epilepsy living in their natural environments. **Top)** Schematic for bi-directional data transmission (using Medtronic Clinician Telemetry Module – CTM) between implanted brain sensing and stimulation device integrated with local handheld computer (Epilepsy Patient Assist Device - EPAD) and cloud environment. The integrated system provides a platform for chronic ambulatory monitoring of patient reported behavior, device data (battery, telemetry, electrode impedances), seizures, and interictal epileptiform spikes (IES). **Bottom left)** The cloud-based Epilepsy Dashboard enables review of electrophysiology data and analytics from a battery of algorithms running on the patient’s local handheld. **Bottom right)** The physician can quickly review and confirm or reject automatically detected and patient reported candidate seizure events. The panel shows 7-days of continuous hippocampal IES rates and seizure detection probability. Blue triangles show patient reported seizure events. Circles denote automated seizure detections either confirmed as seizures (blue dots) or false positive (red) by expert visual review. Zoomed circular inset shows example of raw data from hippocampus with automated IES detections (red circles). The patient was aware and reported (blue triangle) one out of the six seizures detected in the continuous iEEG and confirmed by the physician.

## Materials and Methods

### Study design and data sources

To develop classification algorithms, we used a large database of iEEG from two different implanted devices, the NeuroVista (NV) and Medtronic devices that wirelessly stream iEEG data (Table 1). The development dataset included 13 humans (NV1-9; MH1-4) and 8 canines. The MD1 dog (implanted at Mayo), UCD1 and UCD2 (implanted at UC-Davis), and 5 NeuroVista -dogs with epilepsy implanted across multiple institutions in US^16,17^ had naturally occurring epilepsy. We used 9 NV humans and 8 dogs (MD1, UCD1, UCD2, and 5 NV dogs) for training, validation, and pseudo-prospective testing seizure and IES detection algorithms. Then classification algorithms were prospectively tested within a distributed brain co-processor for neurophysiologic tracking and adaptive stimulation in 2 pet dogs (UCD1, UCD2) and four human subjects (MH1-4) in their natural environments.

**Table 1.** Seizure detection datasets. Training, validation, and testing data used in development of a generic, automated seizure detection algorithm for canines and humans. Retrospective data included human and canine datasets acquired with two different investigational devices, NeuroVista (NV) and RC+S^™^ device. Algorithm training was performed using retrospective data from humans and canines collected with NV devices. The validation data used NV data from two humans (N1, 2) and RC+S^™^ data from three canines (UCD1, 2; MD1). Pseudo-prospective (NV data from 7 humans; NV3-9) and prospective (RC+S^™^ data from 4 patients MH1-4 and 2 pet dogs UCD1, 2) ambulatory testing in human and canines living in natural environments (human at home and dogs living with their owners) was performed over multiple months.

|  |  | training | validation | testing |  |
| --- | --- | --- | --- | --- | --- |
|  |  | Retrospective Data | Retrospective Data | Retrospective Data | Prospective Data |
| NeuroVista Device |  | 5x NV dogs<br>ID (1, 2, 3, 4, 5) |  |  |  |
|  |  | 2x NV Human<br>ID (1, 2)<br>bootstrap 50% of data | 2x NV Human<br>ID (1, 2)<br>bootstrap 50% of data | 7x NV Human<br>ID (3, 4, 5, 6, 7, 8, 9) |  |
| RC+S Device |  |  | 3x RC+S dogs<br>UCD (1, 2)<br>MD 1 |  | 2x RC+S dogs<br>UCD (1, 2) |
|  |  |  |  |  | 4x RC+S Human<br>MH (1, 2, 3, 4) |

#### Devices, training, validation, and testing data

Datasets collected from two implantable devices were utilized for system training, validation, and testing (Table 1). The investigational NeuroVista system is a 16-channel brain sensing (0.1–100 Hz bandwidth; 400 Hz sampling) implantable device providing continuous iEEG wireless streaming to an off-the-body data storage and analytics device carried by the patients and dogs. The RC+S^™^ is a 16 channel electrical stimulation and sensing implantable device capable of selective sensing from any 4 of the 16 channels (1–70, 125, 250 Hz bandwidth; programmable sampling 250, 500, or 1000 Hz) and wireless streaming to a handheld tablet computer with cellular and internet connectivity to a central cloud based data and analytics platform.^11,12^ The investigational NeuroVista and RC+S^™^ devices have yielded massive datasets of ambulatory iEEG in naturalistic settings and are idea for development of robust automated algorithms for brain behavioral state classification, IES and seizure detection. We have previously used the NeuroVista Inc. device data from humans^9^ and canines^18^ for developing seizure detection and forecasting algorithms.^16,19–21^

#### Canine Device Implants

The animal research and clinical care took place at Mayo Clinic, Rochester MN and University of California Davis, Davis, CA under IACUC Protocol A00002655 *Chronic Wireless Electrophysiology and Modulation in Epileptic Dogs*. Epilepsy occurs naturally in dogs with prevalence, age of onset, and clinical presentation similar to human epilepsy.^22^ Naturally occurring canine epilepsy is often drug resistant and new therapies are needed. In addition, the canines provide a platform for preclinical testing, since dogs are large enough to accommodate devices designed for humans. All canines were implanted at either Mayo Clinic (MD1) or at University of California, Davis (UCD1-2).

##### Electrode and RC+S^™^ implantation in dogs

Medtronic deep brain stimulation electrodes were implanted intracranially in canines under anesthesia using a custom-made stereotactic frame. Canines underwent a 3.0T MRI using a stereotactic T1-weighted sequence (Fig. 5). Targets and trajectories were planned using stereotactic software (Compass^™^ Stereotactic Systems) adapted for a large animal head frame. Burr holes were drilled into the skull for each of the four electrodes (Medtronic models 3391 and 3387) that were inserted to the target depth and secured with metal anchors and bone screws. The electrode tails were tunneled to the RC+S^™^ in a pocket behind the canine’s right scapula. The canine underwent a post-op x-ray CT scan, which was then co-registered to the stereotactic MRI (Analyze 12.0, BIR, Mayo Foundation) in order to verify targeting accuracy. We have previously described the similar procedure for the previous NeuroVista Inc. device implants carried out at Mayo Clinic, University of Minnesota, University of Pennsylvania, and University of California Davis in canines.^16,18^

#### Human Subjects

##### NeuroVista device

The 9 human dataset collected with the investigational NeuroVista Inc. device were from the NeuroVista device trial in humans carried out in Melbourne, Australia, between March 24, 2010, and June 21, 2011.^9^

##### Investigational RC+S^™^ Summit

The human subject research with RC+S^™^ was carried out at Mayo Clinic under an FDA IDE: G180224 and Mayo Clinic IRB: 18-005483 “Human Safety and Feasibility Study of Neurophysiologically Based Brain State Tracking and Modulation in Focal Epilepsy”. The study is registered at https://clinicaltrials.gov/ct2/show/NCT03946618. The patients provided written consent in accordance with the IRB and FDA requirements.

We consented 6 patients and implanted 4 patients with drug resistant temporal lobe epilepsy (TLE) as part of the NIH Brain Initiative UH3NS95495 *Neurophysiologically-Based Brain State Tracking & Modulation in Focal Epilepsy*. The details of the approach for implantation have been previously described.^23^ Magnetic resonance imaging was performed after Leksell (Elekta Inc.) frame fixation for stereotactic targeting. Medtronic 3387s electrodes were then implanted in the ANT by direct targeting of the mammillothalamic tract on MRI (FGATIR sequence).^24^ Medtronic 3391 electrodes were implanted into the hippocampus through direct targeting of the amygdala and hippocampal head (Fig. 6.). After confirmation of the electrode location with intraoperative computed tomography (CT), the leads were connected to bifurcated extensions and tunneled to the RC+S^™^ in an infraclavicular pocket. The FDA IDE protocol investigates electrical brain stimulation (EBS) paradigms, including low frequency (2 & 7 Hz) and high frequency (100 & 145 Hz) stimulation, seizure detection and forecasting, behavioral state tracking, and adaptive EBS control.

##### Patient MH1

57-year-old ambidextrous woman with drug resistant mesial temporal lobe epilepsy (mTLE). History of head trauma with loss of consciousness followed by generalized tonic-clonic seizure beginning at age 9. She did well until age 21 yrs., when her seizures became drug resistant. She has comorbid depression and anxiety.

##### Patient MH2

20-year-old right-handed woman with diabetes mellitus type-1 and drug resistant mTLE. No epilepsy risk factors. Epilepsy onset at age 7 years, and a prior left temporal lobectomy at age 9 years. She was seizure free until age 17 years when seizures recurred while off all medications. Thereafter she has been drug resistant. She has comorbid depression and anxiety.

##### Patient MH3

41-year-old right-handed woman with drug resistant mTLE. No clear risk factors for epilepsy. Epilepsy diagnosis was at age 31 years. Despite VNS she had continued seizures. She has comorbid depression and anxiety.

##### Patient MH4

A 35-year-old right-handed woman history of diabetes mellitus and drug resistant temporal lobe epilepsy. She has no epilepsy risk factors. Epilepsy onset at age 4 years old. Significant comorbid depression. Elevated GAD-65 that did not respond to immunotherapy

### Detection of interictal epileptiform spikes

Interictal epileptiform spikes are an electrographic marker of pathologic brain tissue capable of generating unprovoked seizures. In recent years there has been rapid development of new and reliable techniques for automated IES detection. To train and evaluate the IES detector we used continuous hippocampal recordings from the RC+S^™^.^12^ We used a previously validated algorithm^25^ that models and adapts based on statistical distributions of signal envelopes from background (normal) iEEG activity. This enables differentiating signals containing IESs from signals with background activity even in long term data recordings with changing background electrophysiological activity. The IES detector also identified low-amplitude IES in cases where the background activity power is low and IES are often missed by expert visual review.

We benchmarked the IES detector using data acquired with a chronically implanted brain stimulator. We deployed the detector in a cloud system that received the continuously streaming hippocampal data over one year. We compared the detector performance with the manual visual review (GW & NG electroencephalographers) scoring in selected epochs (see Data for IES Detector). The IES detector ran during different anterior nucleus of the thalamus (ANT) stimulation paradigms, no stimulation, 2, 7, and 145 Hz stimulation) with changing stimulation current amplitudes (2, 3, 5 mA) and pulse widths of 90 and 200 usec.

To investigate how IESs characteristics change in periods of different seizure frequency we selected epochs of the data in periods of frequent (cluster) and less frequent seizure activity (non-cluster). The seizure cluster period was defined as more than two seizures in a day. For each of the two (cluster, non-cluster) we selected 5-minute-long epochs for left and right hippocampal channels. Each selected epoch was taken at distinct times to assess differences between sleep and wake cycles. In total we selected twenty-four 5-minutes long epochs reviewed independently by two electroencephalographers. All IESs were marked in both hippocampal channels and used subsequently to calculate congruence score between experts and to validate the automated IES detector. Subsequently, we used the two-months period of the data continuously streamed from the human with implanted RC+S^™^ to analyze IES rates and IES characteristics.

### Generic seizure detector

The training dataset consists of long term NeuroVista recordings from 5 canines and 2 human patients (Table 1). In canines, all seizures were included in training (340 in total). Another 628 interictal segments with various electrophysiological activity patterns were manually selected. The human dataset consists of 1049 seizures and 846 interictal segments. Half of the seizures (524) and half of the interictal segments (423) were bootstrapped and used as training data and the other half of data used in the validation dataset. The validation dataset included two sets of data. The first dataset has data from RC+S^™^ recordings from three canines. Each recording spans at least 210 days. In total, 133 electrographic seizures and 833 interictal segments were selected from the continuous recordings upon visual review by an expert reviewer. The second dataset contains the other half of the data (2 NeuroVista patient recordings) generated by bootstrapping in the training dataset.

The testing datasets include previously collected NV datasets that were used for pseudo-prospective testing and RC+S^™^ datasets for prospective testing. The pseudo-prospective NeuroVista human data was from 7 patients and four human subjects, and two pet canines implanted with the RC+S^™^ system. The pseudo-prospective NV human dataset period is ∼10.5 years and includes 2046 seizures in total (Table 2).

**Table 2.**
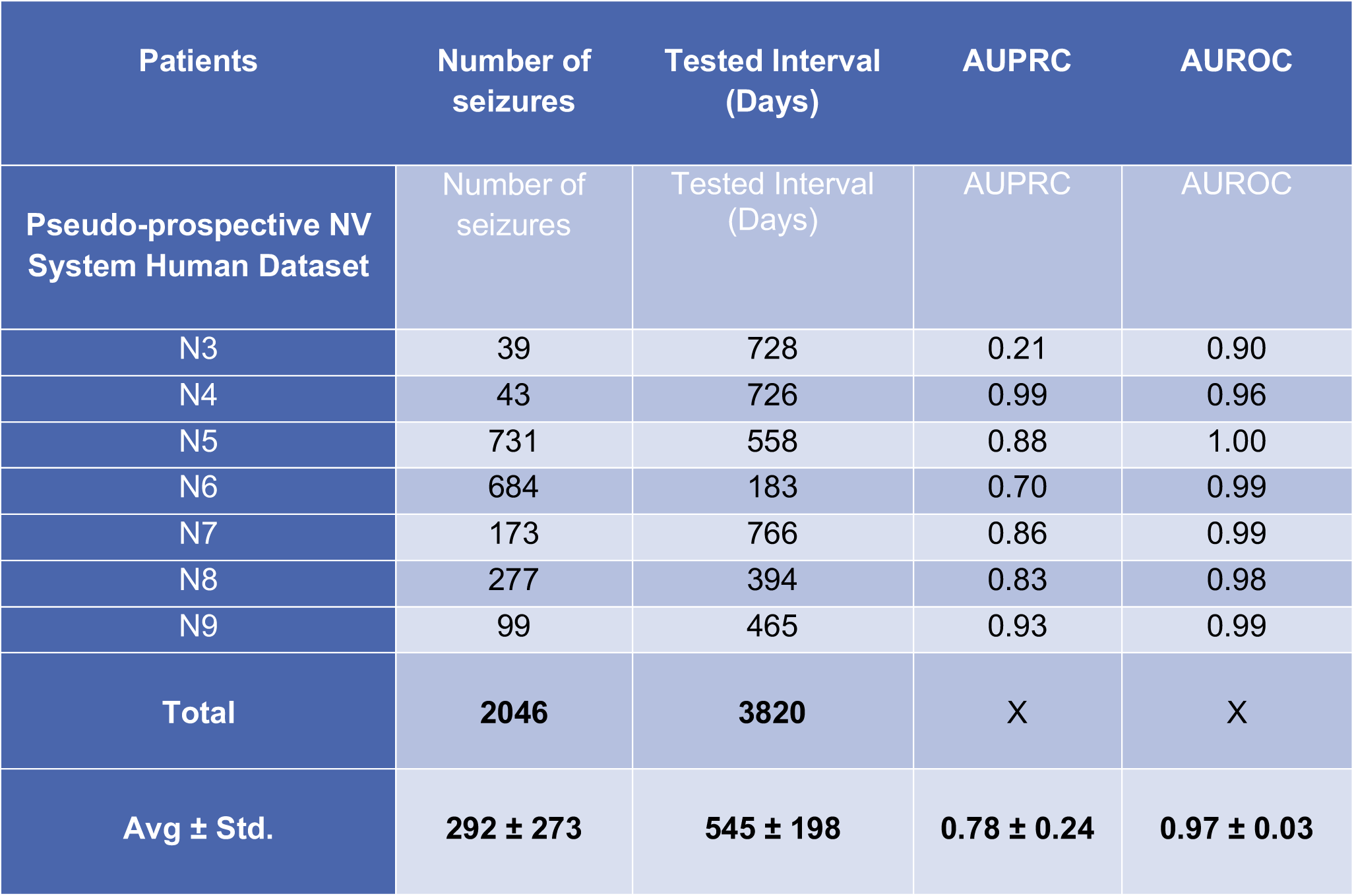
Seizure detection results using NeuroVista Human Dataset. Performance of the generic seizure detection model for canine and human seizures deployed on out-of-sample human NV dataset in pseudo-prospective testing. Pseudo-prospective data was previously collected but analyzed while maintaining the temporal relationship of all seizures. Machine learning performance metrics are shown together with the number of seizures and number of recording days in the datasets (AUPRC - area under precision recall curve; AUROC - area under receiver operating characteristic curve).

The prospective deployment ran over 723 days and contains 204 seizures that were recorded in the four humans (MH1-4) and two pet canines (UCD1 and UCD2 RC+S^™^ system).

#### Detector Design - utilizing LSTM neural network

To design a generalizable seizure detection algorithm for a generic implantable system, we required the algorithm operate independently of the recording system, spatial electrode position, and species tested. We used two of the few implantable neuro-devices capable of continuous streaming local field potential data through a wireless connection. This allows long-term, real-time monitoring since the collected data are continually transferred from the implantable device to the brain co-processer system (tablet or smartphone, and cloud computational resource) ^12^. For this reason, the algorithm must be capable of processing data streams with artifacts and data drops caused by interference or disconnections.

Seizures recorded with intracranially implanted electrodes exhibit temporal evolution of spectral power across a wide range of frequencies. Different electrographic signatures are observed in the data based on their initial power distribution. It is important to note that seizures in one patient might have multiple ictal patterns, therefore training on different ictal patterns is necessary for high sensitivity seizure detection. The detector has to distinguish ictal patterns from sharp transient artifacts coming from a recording device or short interictal discharges which might temporarily increase spectral power similar to an electrographic seizure.

Previously reported seizure detectors^20,26–28^ usually utilize combination of features extracted from multiple channels, or features extracted from shorter segments without adaptation to a long-term baseline. This is a crucial design input requirement in designing the seizure detector for a long-term monitoring in chronically implanted devices. Another drawback of previously reported detectors is that the testing is usually done on isolated ictal and interictal segments, and not on a long-term continuous recording spanning weeks and months of time. The deployment of trained and validated seizure detectors on previously unseen out of sample unbalanced data is critical for evaluation and real-time performance of a generalizable seizure detector. Our method focuses on spectral changes in iEEG recordings from only one channel and marks a probability of seizure over time thus providing independence from the neighboring channels and the short interictal discharges that could confuse current state of the art detectors.

In order to address these requirements, we developed a convolutional long short-term memory (LSTM)^29,30^ neural network utilizing Short Time Fourier Transform (STFT) calculated from single lead iEEG as an input. We previously used CNN with LSTM for automated classification of iEEG.^31^ The STFT converts the single lead time series data into time-frequency representation (spectrogram). Invariance to sampling frequency is achieved by using a constant time window of 1 second with 0.5 seconds overlap, and subsequently selecting only frequencies lower than 100 Hz. A raw data segment is always transformed into a spectrogram image with dimensions 100xT, where T is the number fast Fourier transform (FFT) calculations, not depending on sampling frequency (frequency domain resolution is always 1Hz per sample). Time series data of 5 min length were empirically chosen to provide long enough EEG baseline temporal context for the LSTM, so the relative power of seizure stands out of the background activity. The final classification is made for every 0.5 sec of the 5-minute input raw data signal using a many-to-many LSTM architecture. Raw data are z-score normalized prior to STFT calculation and each frequency band of the resulting spectrogram is z-score normalized prior to the neural network inference. Dropout layers in neural networks are used for regularization during training to prevent overfitting. Similarly, we drop random segments prior to the spectrogram computation. This enables the network to handle the data from the wireless system with possible short data gaps.

The convolutional LSTM model consists of 2 convolutional blocks (convolution and ReLU) with kernels {5, 5} and {96, 3}, respectively. Subsequently, time distributed feature representation is processed with 2 layers of bidirectional LSTM recurrent neural network. Lastly, a fully connected layer with a softmax activation function transforms the LSTM output into probability output. The proposed architecture is trained with Adam optimizer (learning rate = 10^−3^, weight regularization = 10^−4^) in a many-to-many training scheme, where every input FFT window has a multiclass label. We implemented 4 types of labels – normal activity, IES together with artifacts, dropout segments, and seizures. Adding additional labels might improve learning because the model is forced to not only distinguish interictal activity from continuous seizure activity but also interictal discharges which are not considered as electrographic seizures in different behavioral states, and thus lower the number of false positives. The temporal resolution of the detector is defined by the FFT window step (0.5 seconds). In order to train the network, we use a special purpose deep learning computer Lambda Labs Inc. (8x GTX 2080TI GPU, 64 CPU cores and 512 GB RAM). The data-parallel training method runs on all GPUs and average model gradients and is used to reduce training time. The model is built in the PyTorch deep-learning library for Python.

#### Training and validation of seizure detection model

The model was trained on NeuroVista data (5 canines, 2 human patients, Table 2). All training segments were 10 minutes long. Random 5 minutes intervals were sampled from the full segments during the training every time the segment was used in training. Because the human training dataset had a higher number of examples than the canine training dataset during the training epoch training examples were randomly sampled in a way that the number of examples from both classes was balanced.

Performance of the model during the training was evaluated by area under the precision-recall curve (AUPRC), where all seizure targets were set to one and all the other classes were set to zero. Validation of CNNs is typically measured by validation loss, but we used AUPRC for scoring because it is independent of the probability threshold of the classifier and it is not dependent on the true negative samples in the dataset. Validation examples were fixed 5-minute intervals and were not randomly sampled. Validation scores (AUPRC) were calculated on two different datasets (3 canines with RC+S^™^, 2 human patients with NeuroVista device) independently. The two validation scores were averaged after each training epoch and the model with the best score achieved during training was deployed on the test dataset in order to obtain results (Table 2).

#### Model deployment

We arbitrarily chose 10 continuous seconds of ictal activity as an electrographic event that we want to detect ^32^. The model iterates over the data with 5-minute windows with 100 seconds of overlap. The model gives a probability of seizure for every 0.5 seconds (higher probability is used in the overlap region) in every channel. Seizures in the test dataset are marked across all channels without specification therefore we combine probabilities from all channels in the following way. The three highest probabilities from all channels are averaged and from this averaged probability the final performance measures are calculated. For a given probability threshold the system identified continuous detection whenever the probability was above a threshold (see example of a detection in Fig. 2). Next, every detection interval above a threshold was automatically extended if in the next 10 seconds from the current detection was another detection. Subsequently, the two detections were merged into one interval. Thus, for every probability threshold, we detected intervals of various lengths which the model marks as seizures. Intervals shorter than 10 seconds were dropped from detected events. For detected events longer than 10 seconds AUPRC and area under the receiver operator curve (AUROC) scores were calculated based on the region overlap with gold standard seizures marked by an expert reviewer.

**Fig. 2.**
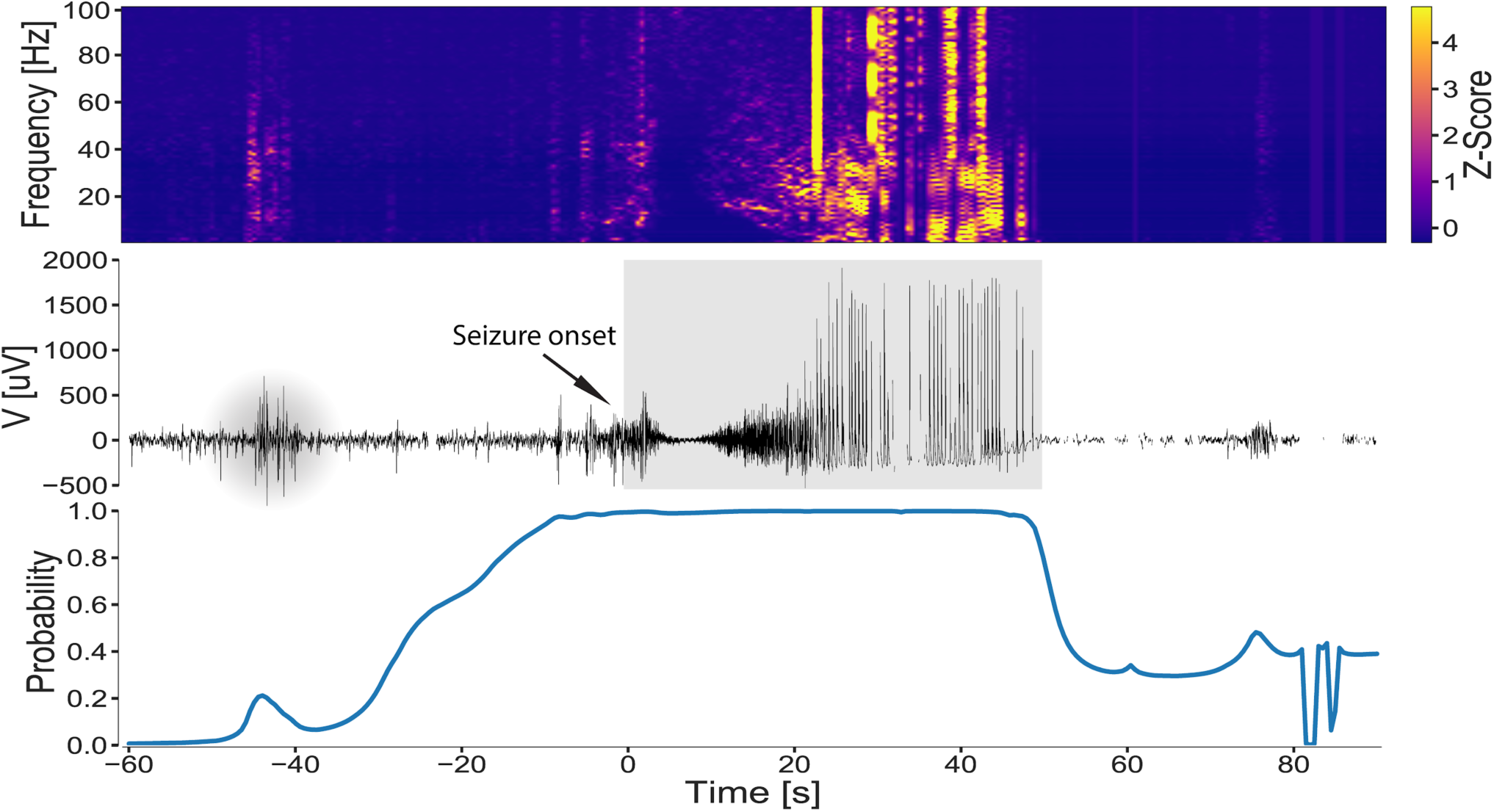
A representative hippocampal seizure from human subject MH1. **Top)** Time-frequency characteristics (z-score spectrogram), **Middle)** Raw intracranial EEG data with the physician annotated (grey highlight) seizure duration, and **Bottom)** Model seizure probability for patient MH1 in the out-of-sample dataset demonstrating how the probability of the long-short term memory (LSTM) model changes over a peri-seizure period (pre-ictal, ictal, and post-ictal period). The high probability (near 1) in the peri-seizure region highlights the impact of the LSTM function for raising the probability during and around the seizure time.

The model was deployed in local cloud storage to continuously process incoming data from RC+S^™^ animal and human study. Due to a different electrode configuration in the RC+S^™^ system in comparison with the NeuroVista system, we could not use an average of the three highest probabilities. Instead, a maximal probability given by two hippocampal channels was taken as an output of the model. Subsequently, the detected intervals were calculated from the probabilities in the same manner as for the data from the NeuroVista dataset. The model has been running online and continuously detecting seizure events as the new data were coming in. A revision of the raw data by an expert reviewer created gold standard seizure marks for comparison of classifier performance. Thus, with all detected events and true seizure marks AUPRC and AUROC scores were calculated.

The performance of the model on out-of-sample data is numerically shown in Table 2. The performance of the generalized classifier is visualized using standard machine learning graphs of PRC and ROC for each individual human (Fig. 3). The results of model detections outperform state of the art detectors published recently Baldassano, Brinkmann et. al^20^ and directly compare two hundred teams of data scientists across the globe comprising 241 individuals. An advantage over a Kaggle competition we were able to take and use a larger portion of the full dataset in a more realistic setting, where the classifier is trained and then pseudo-prospectively run on the new out-of-sample data of different subjects in a sequential way fully simulating a real prospective situation of recording where new data are arriving each second and detector runs in near real-time manner. Yet, the classifier doesn’t need to be retrained for patient specific applications and is fully generalized. On the other hand, Fig. 7 shows an example of a short period (a minute) of iEEG data with seizure for all sixteen neocortical electrodes of patient NV7 from NV human dataset. The seizure is visually apparent in only a few channels with adequate signal to background ratio suitable for automated detections. This is likely a common situation with electrodes spanning the space from seizure onset zone to surrounding regions of the brain. Fig. 7 reveals the time-frequency analysis of these iEEG signals showing the different signatures of seizure electrophysiology in different channels and below is the visualization of the classifier output probabilities for each electrode. This also shows in the time-frequency domain that for some electrodes the seizure is very prominent while for others not differentiable from the background signal. Therefore, here the model decides based on the seizure probabilities of the electrodes taken as a mean of top three probabilities.

**Fig. 3.**
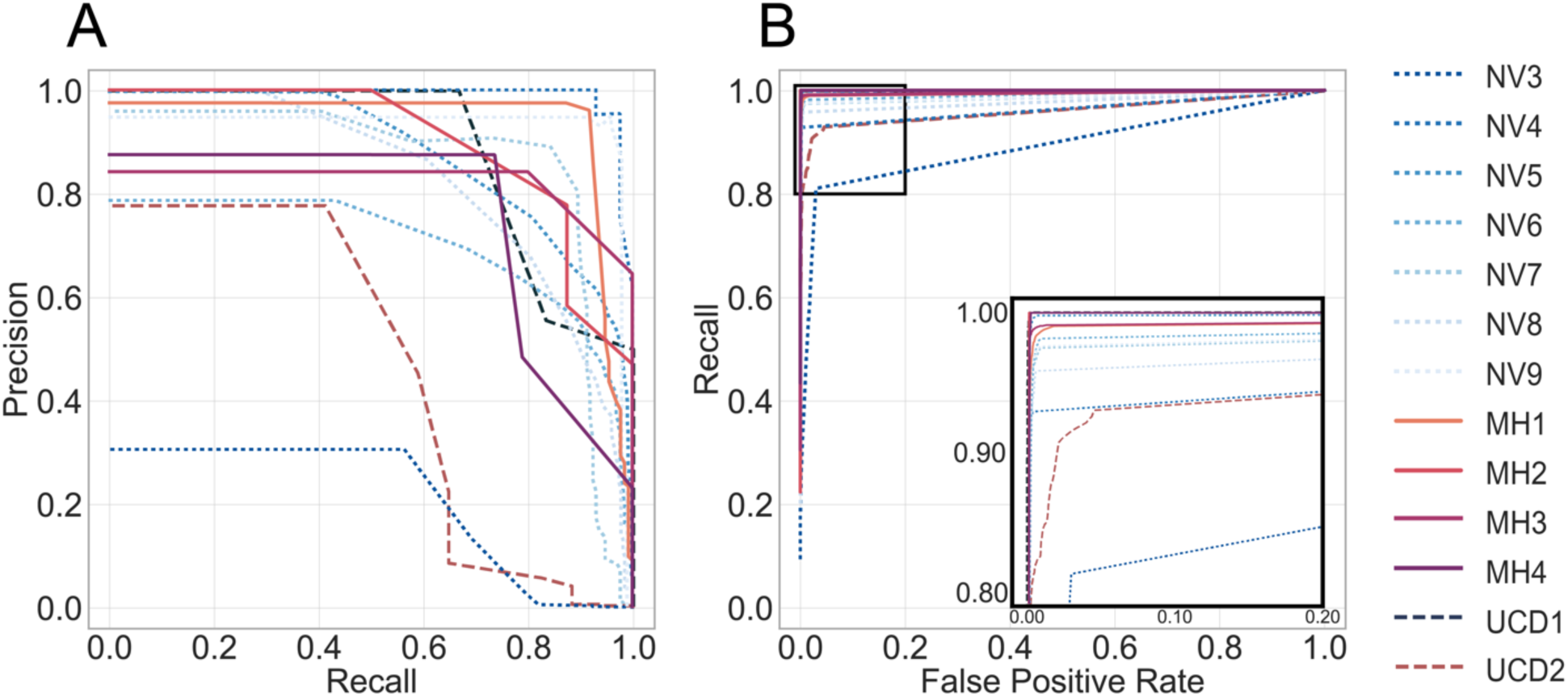
The long short-term memory model performance. Out-of-sample retrospective testing in human (dotted NV3-9), and prospective testing in human (solid lines MH1-4) and canine (dashed lines) subjects: **(A)** Precision Recall Curves (PRC) and **(B)** Receiver Operating Curves (ROC). The detailed view of the ROC (blow-up view in bottom right panel) shows the results for each subject with optimal detector parameters that minimize the false positive rate and maximize sensitivity. The PRC and ROC curves are calculated by sequentially changing the model probability threshold and evaluating the results of Precision, Recall, and False Positive Rate for all seizures for each subject in the testing datasets.

##### Data and materials availability

All results associated with this study are present in the paper. The data and analysis code are available upon reasonable request (https://www.mayo.edu/research/labs/bioelectronics-neurophysiology-engineering/overview)

## Results

### Tracking behavior and epilepsy biomarkers in humans & canines

We used analysis of intracranially recorded electroencephalography to detect seizures and IES in ambulatory humans and canines with drug resistant epilepsy living in their natural environments. Continuous streaming iEEG was analyzed in a cloud environment and on a tablet computer, carried by subjects that also enabled synchronized patient inputs. Physicians and engineers remain in the loop using a web-based Epilepsy Dashboard to review biomarker trends (IES rates & seizures), patient annotations (seizures, auras, medication logs), and implanted device data (battery status, telemetry, and EBS parameters). The system provides an integrated machine learning platform for algorithm development, data viewing, biomarker tracking and expert annotation of events, e.g. confirmation that a detected electrophysiological event or patient reported event was a true positive seizure (Fig. 1).

### Automated seizure detection

Accurate seizure catalogues are critical for optimal epilepsy management and assessment of EBS outcomes, but remain a basic technology gap for the field.^6–8^ We created an accurate seizure diary based on a generic seizure detector using a Long-Short-Term-Memory (LSTM)^30^ artificial recurrent neural network (RNN) and convolutional neural network (CNN)^29^ applied to continuous iEEG to reliably detect seizures in ambulatory canine and human subjects with epilepsy.

The large testing, validation, and training dataset from multiple brain structures in humans and canines was collected over multiple years with two different fully implantable recording devices (NeuroVista Inc. or Medtronic PLC, see methods). The LSTM model was trained on a dataset from five dogs with naturally occurring epilepsy (340 seizures) implanted with NeuroVista devices^16,18^ and one half of the data from two, randomly selected human subjects with epilepsy implanted with the NeuroVista device (524 seizures). The model was then validated on the other half of the data from two NeuroVista patients (524 seizures) and three canines (133 seizures in dogs: UCD1, UCD2, MD1) implanted with RC+S^™^ devices (Table 1).^12^

Automated detection of spontaneous seizures recorded with iEEG is possible because of the characteristic spectral patterns that are readily identified visually and by machine learning approaches.^17^ Figure 2 shows an example of a typical seizure with its time-frequency (spectrogram) characteristics, raw data, and LSTM model seizure probability for MH1 from the out-of-sample data. The LSTM model probability for seizure classification changes in context of raw iEEG and spectral content showing high probability within the seizure activity and low probability outside the seizure (before and after the seizure). The example highlights the importance of the LSTM in the model, since feature-based machine learning models would detect the bursts of IES at the beginning and during the seizure, while the LSTM model raises the seizure probability prior and during the seizure time.

The precision recall curves (PRC), and receiver operator curves (ROC) curves are calculated by sequentially changing the model probability detection threshold and evaluating the results for all seizures from each subject in the testing datasets (Fig. 3).

The performance of the generalized automated seizure detector using out-of-sample data from 7 human patients implanted with the NeuroVista device (total seizures 2046 over 3820 days) was AUPRC 0.78 ± 0.24 and AUROC 0.97 ± 0.03 (Table 2).

### Detection of interictal epileptiform spikes

Interictal epileptiform spikes are an established biomarker of epileptogenic brain,^33^ and associated with risk for spontaneous, unprovoked seizures.^34–36^ For long iEEG datasets it is labor intensive and impractical to use visual analysis to calculate IES rates. Here we trained, validated, and tested an automated IES detector on long-term continuous ambulatory iEEG recordings. We implemented a previously published automated IES detection algorithm,^25^ where the data are continuously accumulated by streaming iEEG from the RC+S^™^ device to a cloud database. We compared the automated IES detections to expert visual scoring from two epileptologists (NG & GW). These data included periods during day, night, seizure clusters (2 or more seizures in 12 hours) and non-seizure cluster periods. There was good concordance for the IES labeling by expert visual review (Cohen’s kappa score 0.87) and between the algorithm and experts (F1-score 0.82 ± 0.08 with sensitivity 91 ± 0.6% and positive predictive value 77 ± 1.6%).

The algorithm performs well during night, day, high and low seizure periods (Table 3). The IES rates are higher during seizure clusters periods (2 or more seizures in 12 hour period), but performance of the automated detector is similar during periods with high and low IES rates (F1-score was 0.84 in seizure cluster and 0.80 in non-cluster seizure periods).^36^ Despite the difference in IES rates between day (approximately 25% lower IES rates) and night the algorithm performed similarly (day F1-score was 0.81 and 0.82 at night). Visual examples of IES and comparison of automated detections with expert visual review are shown in Fig. 4 for day (A) and the night (B) and illustrate the concordance between expert visual review and the automated classifier. The hippocampus IES rate variations during day and night over a two-month period show circadian and multi-day fluctuations (Fig. 4C). We analyzed IES characteristics to explore how the hippocampal IES properties differ in various behavioral states (Fig. 4D) and find higher peak to trough IES amplitudes during night compared to wake for all four human subjects (p<0.001).

**Table 3.**
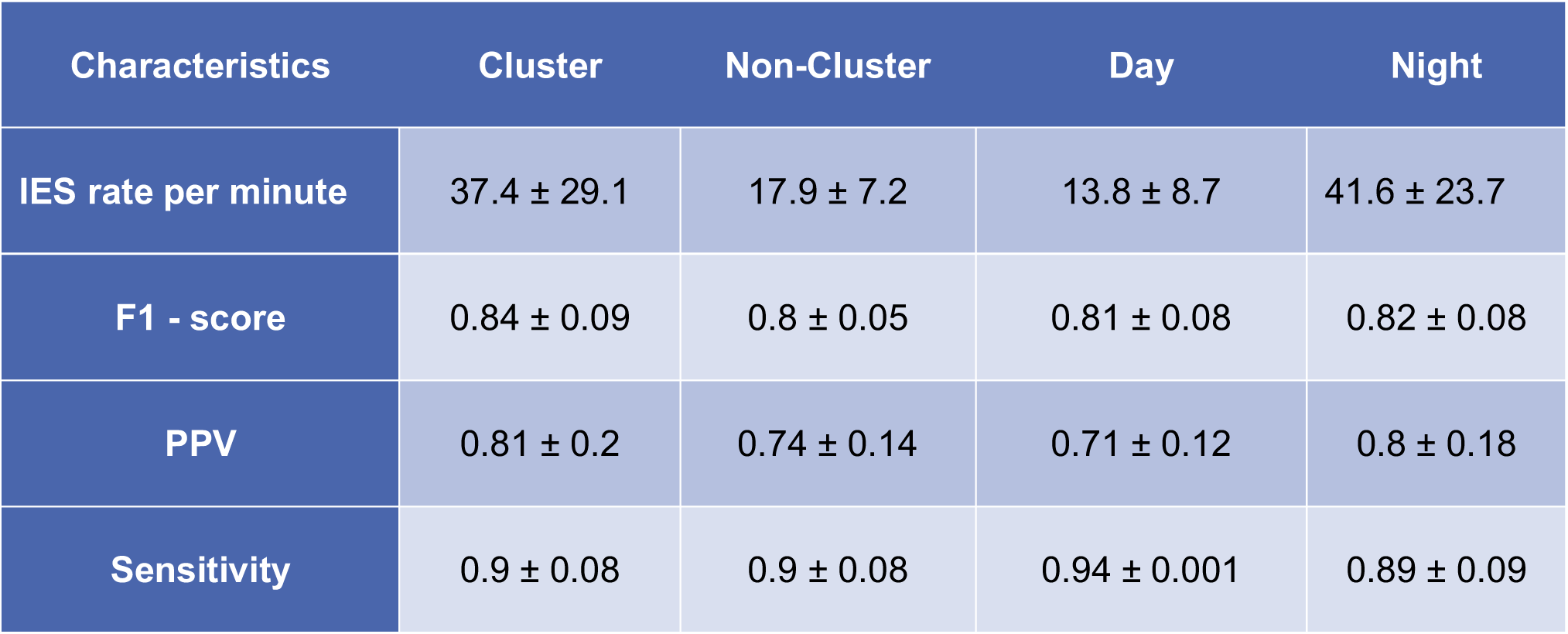
Interictal epileptiform spike rates. Results from prospective testing of the automated IES classifier at different time periods (Day vs. Night) and seizure counts (seizure clusters/non-clusters) compared to expert visual review. Periods of seizure clusters were defined by two, or more, seizures in a 12-hour period. The F1-score comparing the automated detector and expert visual review for labeling IES was similar for each condition studied.

| Characteristics | Cluster | Non-Cluster | Day | Night |
| --- | --- | --- | --- | --- |
| IES rate per minute | 37.4 ± 29.1 | 17.9 ± 7.2 | 13.8 ± 8.7 | 41.6 ± 23.7 |
| F1 - score | 0.84 ± 0.09 | 0.8 ± 0.05 | 0.81 ± 0.08 | 0.82 ± 0.08 |
| PPV | 0.81 ± 0.2 | 0.74 ± 0.14 | 0.71 ± 0.12 | 0.8 ± 0.18 |
| Sensitivity | 0.9 ± 0.08 | 0.9 ± 0.08 | 0.94 ± 0.001 | 0.89 ± 0.09 |

**Fig. 4.**
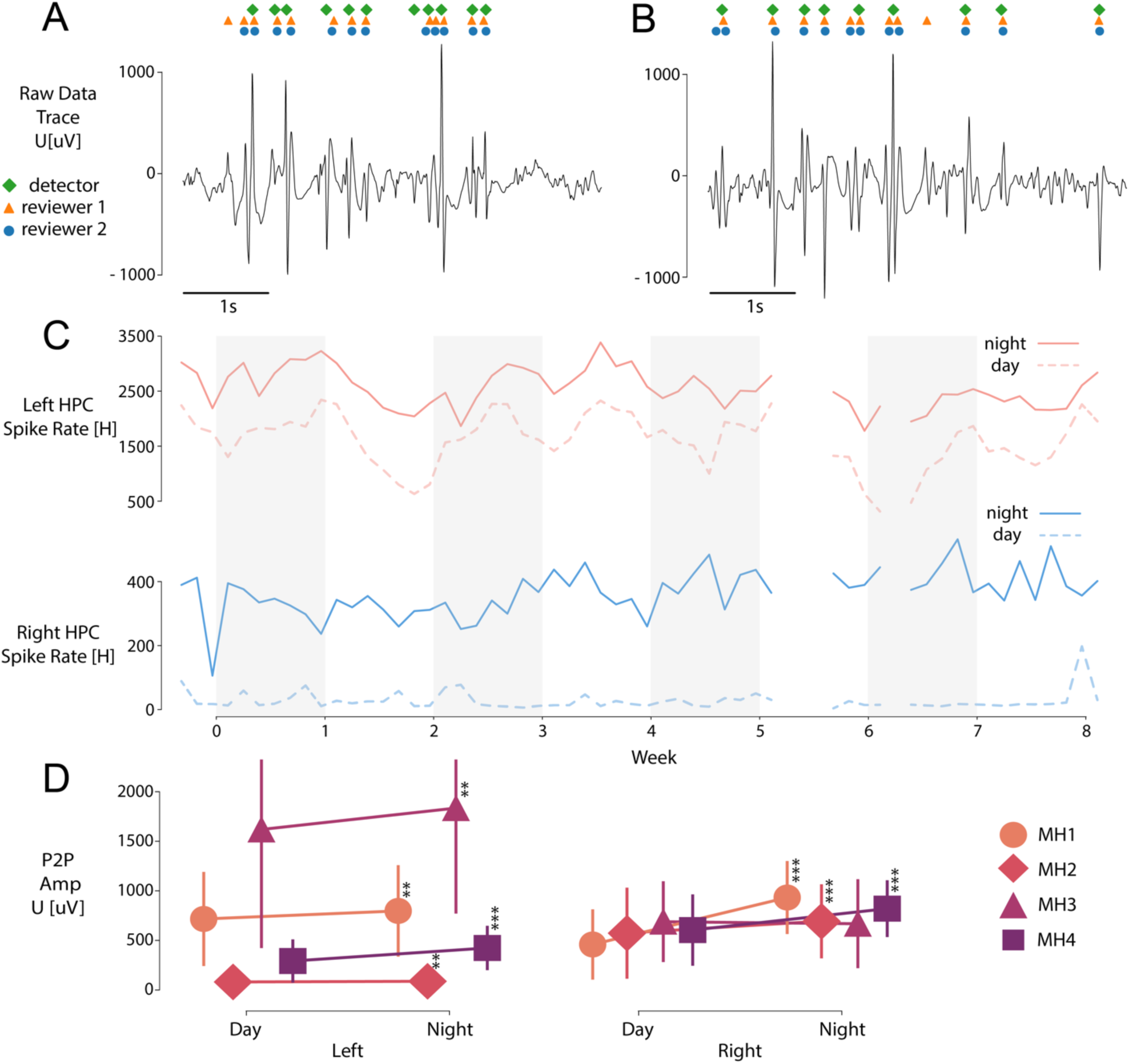
Long-term analysis of Interictal Epileptiform Spike (IES) rates. Visual example of comparing spike detections between the automated approach and human operators for in **(A)** day/awake and **(B)** night/sleep period. **(C)** Daily averaged spike rate per hour in left (top) and right (bottom) hippocampus during night and day periods of time in eight weeks of MH1 recording. **(D)** There are significant differences between night/day in left hippocampal IES peak-to-peak amplitudes during the prospective testing period for all four patients implanted with RC+S^™^.

**Fig. 5.**
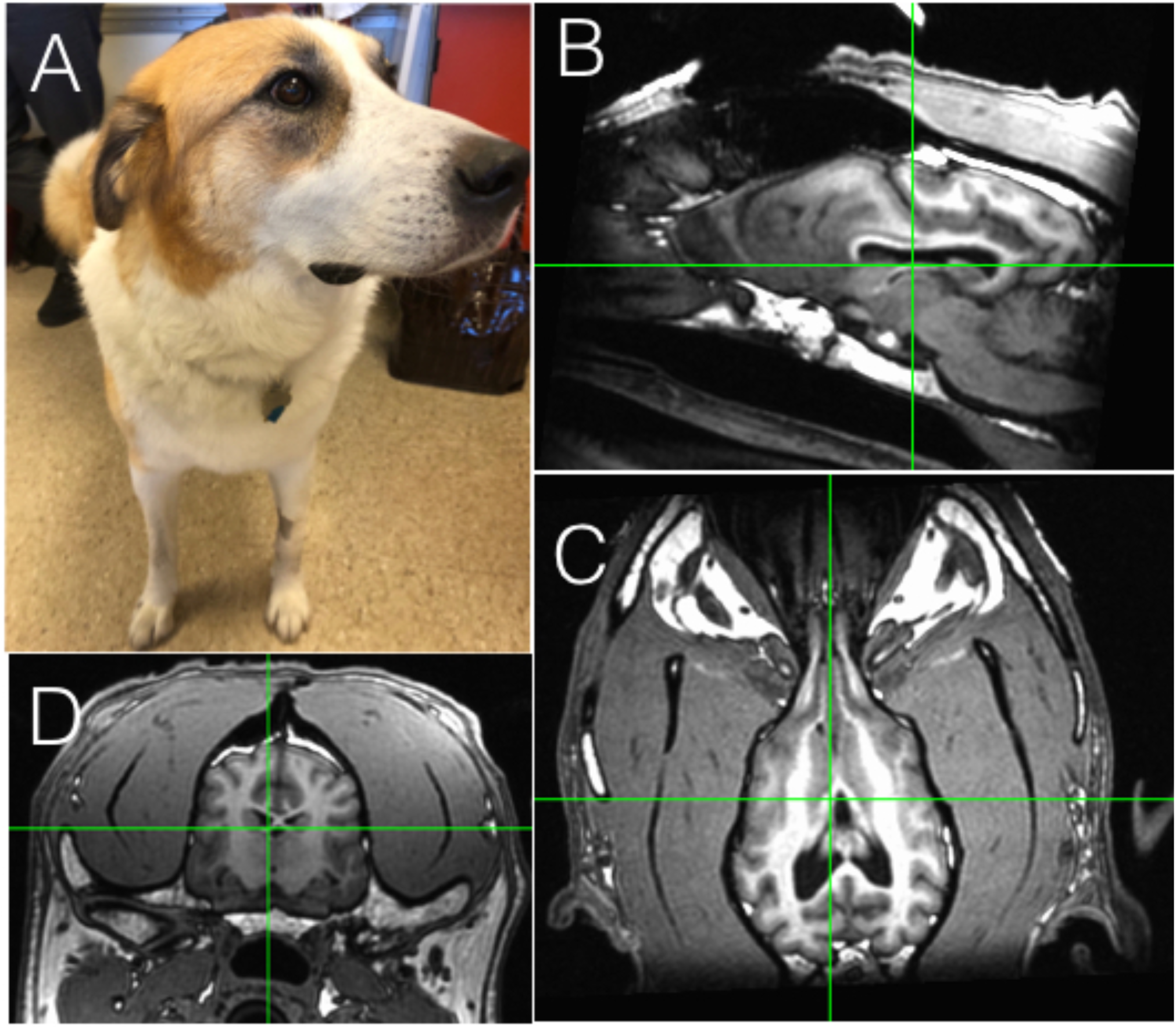
Canine stereotactic implant. **(A)** 6 yr. old pet dog with drug resistant epilepsy. High resolution **(B)** Sagittal. **(C)** Axial. **(D)** Coronal T1 MRI. The electrode implants are by direct visual targeting of anterior nucleus of thalamus and hippocampus.

**Fig. 6.**
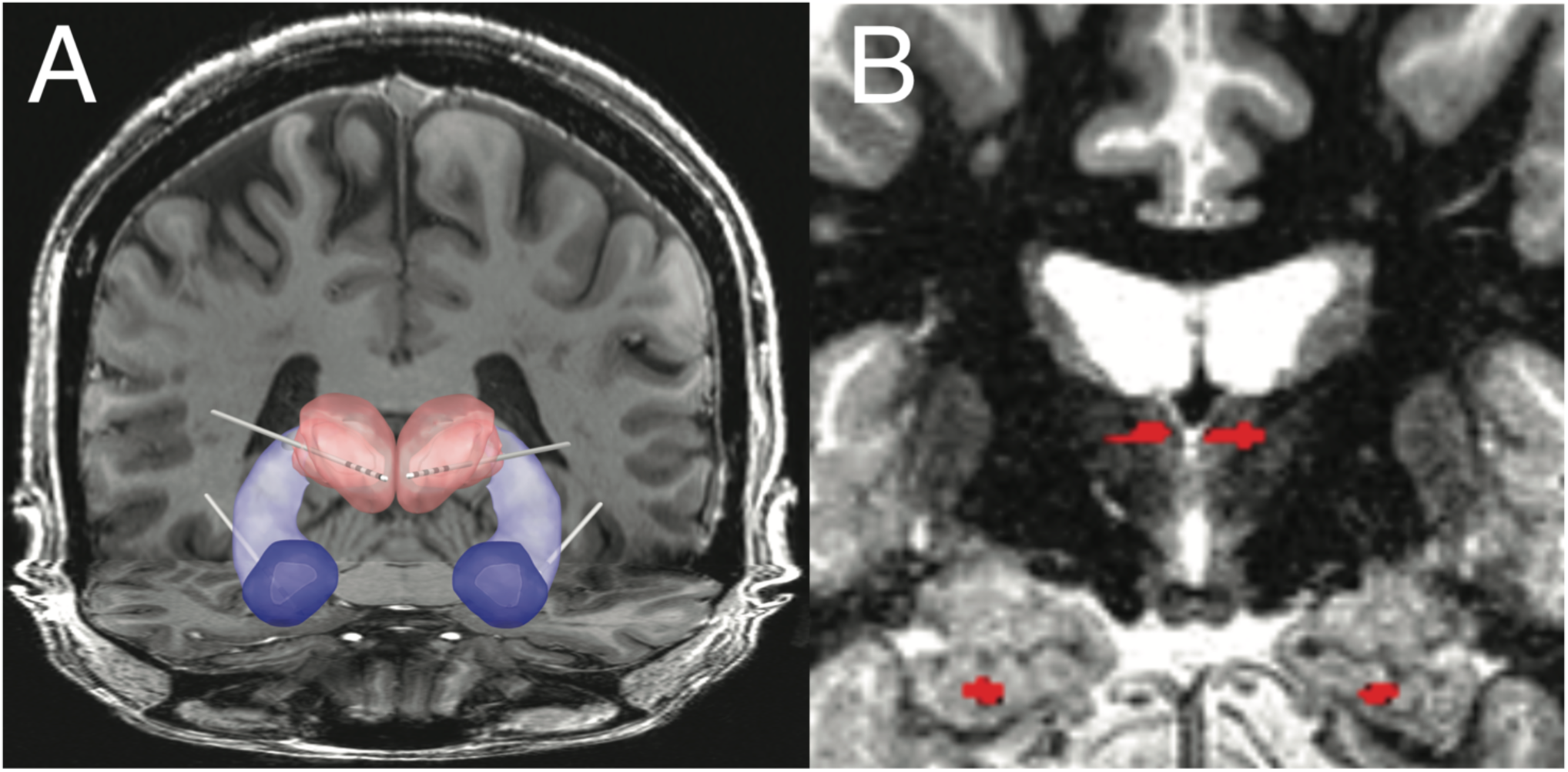
Human subject MH1. (A) Bilateral Anterior Nucleus Thalamus (ANT: red) and Hippocampus (HC: purple) and amygdala (AMG: blue) implant. Papez circuit and implanted electrodes. (B) MRI - the ANT and HC electrodes from co-registration of MRI and post-implant CT are highlighted in red.

**Fig. 7.**
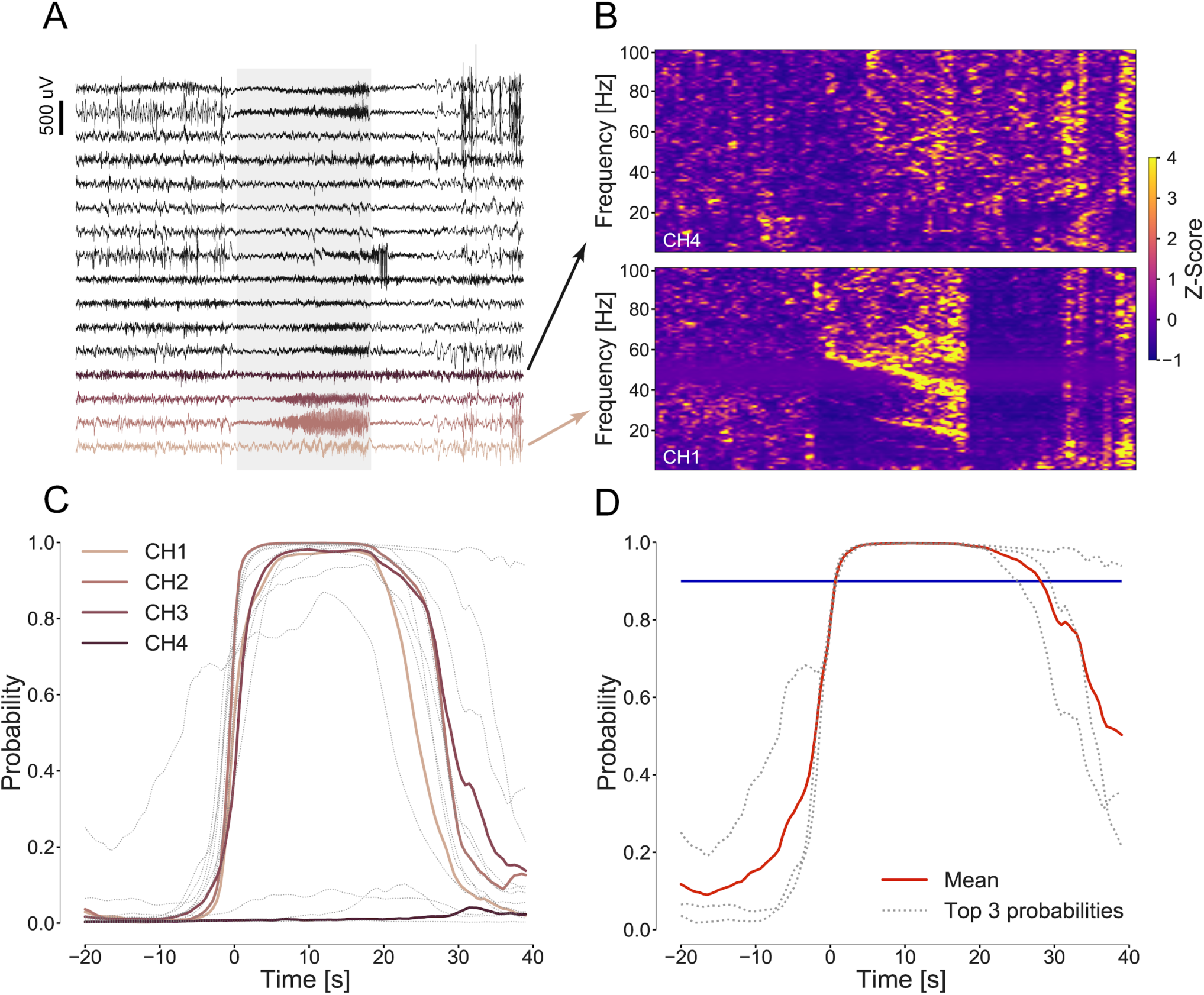
Spectral features of spontaneous seizures. (A) One minute of iEEG data recorded with NV device, sixteen neocortical electrodes, containing a spontaneous human seizure. The seizure is present on a few channels with a good signal to background ratio suitable for automated detection. (B) Time-frequency analysis of signals show the different signatures of seizure electrophysiology (shaded area) in different channels: channel 1, where seizure is notable and channel number 4 where it is hard to identify the seizure. (C) Plots of classifier probabilities for each electrode below actual raw data showing that for some electrodes the seizure is very prominent and for some not differentiable from the background signal. (D) The classifier output probabilities for top three probabilities together with the mean (red) and threshold (blue) showing when the detection is raised (time 0).

### Prospective long-term ambulatory monitoring and algorithm testing

After training, validation, and retrospective testing using previously collected data we then deployed the automated IES and seizure classifiers prospectively in four humans (subjects MH1-4) and two pet dogs (UCD1-2) with epilepsy living in their home environments. In total, the system was able to record an average of 66 ± 0.17% of the data across all human subjects.

The performance of the IES detection in the ambulatory prospective data compared to gold standard expert visual reviewed events was 0.90 sensitivity and F1-score of 0.81. (Table 3).

The prospective testing of the seizure detector in ambulatory subjects in real-world environments showed excellent performance, with an area under the ROC of 0.99 ± 0.01 and PRC of 0.76 ± 0.25 using the expert visual review of the continuously acquired iEEG as the gold standard for the humans (MH1-4) and the two pet canines (UCD1,2). The area under the PRC that more accurately describes the results of this highly imbalanced data with over 99% of the time spent in a non-seizure (interictal) state was 0.93, 0.89, 0.82, 0.75 and 0.47, 0.88 for the four humans and two pet canines (UCD1,2), respectively (Table 4a).

**Table 4a.** Prospective seizure detection results. Performance of automated seizure detection in canine and human seizures deployed prospectively in pet canines and humans living with epilepsy in their home environments (MH1-4 are four human subjects and UCD1-2 are the two pet dogs). Machine learning performance metrics are shown together with the number of seizures and number of recording days in the datasets (AUPRC - area under precision recall curve; AUROC - area under receiver operating characteristic curve).

| Prospective RC+S<br>Summit System Dataset | Number of<br>seizures | Tested Interval<br>(Days) | AUPRC | AUROC |
| --- | --- | --- | --- | --- |
| MH1 | 134 | 147 | 0.93 | 0.99 |
| MH2 | 8 | 149 | 0.89 | 0.99 |
| MH3 | 20 | 44 | 0.82 | 0.99 |
| MH4 | 19 | 156 | 0.75 | 0.99 |
| UCD1 | 17 | 107 | 0.47 | 0.96 |
| UCD2 | 6 | 120 | 0.88 | 0.99 |
| **Total** | **204** | **723** | X | X |
| **Average ± Std.** | **54 ± 49.34** | **120.5 ± 41.97** | **0.76 ± 0.25** | **0.99 ± 0.01** |

The human subjects reported a total of 555 seizures using the EPAD over the course of 945 days of monitoring, but only 39.71 ± 29.20 % were associated with an electrographic correlate (verified seizures) (Table 4b).

Interestingly, of the 407 detected electrographic (ECoG) seizures 43.86 ± 30.77 % were not identified by the patient (Table 4b).

**Table 4b.** Analysis of patient seizure diaries and electrographic seizures from continuous electrocorticography. Continuous invasive electrocorticography (ECoG) from bilateral amygdala, hippocampus and anterior nucleus of thalamus enabled direct assessment of patient seizure diary reports and ECoG recorded electrographic seizure activity. On average 39.7% of ECoG electrographic seizures had a patient seizure diary report within ±30min of the ECoG event and 56.14% of the patient reported seizures had associated ECoG confirmed events. These results from continuous ambulatory ECoG in natural environments demonstrate the complex, and inaccurate relationship between patient diary reports and gold-standard continuous ECoG. *Note:* All four patients had seizures independently from left and right hippocampus. The table includes all seizures, except for MH2 where the results are for right-hippocampal seizures only because of very frequent electrographic seizures (average 30/day) originating from the left hippocampal remanent of a prior left anterior temporal lobectomy. **MH4 excellent reporting included her caregiver reports. The accuracy of patient reporting of seizures is currently confounded by fact that ECoG seizures may be truly subclinical seizures or amnestic seizures that the patient does not recall.

| Subject | Days monitored | ECoG Seizures | Reported Seizures | Patient Seizure Reports & ECoG |  |
| --- | --- | --- | --- | --- | --- |
|  |  |  |  | Patient report with ECoG seizure [%] | ECoG seizures not reported by patient [%] |
| MH1 | 269 | 273 | 80 | 38.75 | 88.64 |
| MH2* | 258 | 13 | 279 | 2.87 | 38.46 |
| MH3 | 153 | 89 | 165 | 43.03 | 20.22 |
| MH4** | 265 | 32 | 31 | 74.19 | 28.12 |
| **Total** | **945** | **407** | **555** | X | X |
| **Avg±Std.** | **378 ± 287** | **163 ± 153** | **222 ± 187** | **39.71 ± 29.20** | **43.86 ± 30.77** |

## Discussion

There has been significant progress in EBS devices for drug resistant epilepsy, but the time to achieve optimal individualized stimulation parameters is long and complete seizure free outcomes remain relatively rare. We suspect that the ability to continuously track electrophysiology, seizure counts, and patient behavior will accelerate the optimization of individualized EBS therapy. To address the technology gaps in currently available EBS systems we developed and deployed a distributed brain co-processor to investigate and continuously track patient reported symptoms, IES biomarkers, and seizures during EBS.

We show that seizures and hippocampal IES rates and characteristics are dynamically changing in a circadian pattern with IES rates highest at night. The hippocampus IES rate variations showed a circadian fluctuations and higher peak to trough IES amplitudes during the night. The accurate automated quantification of IES is potentially of fundamental importance in epilepsy.^37,38^ Interestingly, seizures preferentially occurred during wakefulness in the human subjects despite increased IES rates during sleep.^39,40^ The reason for this phenomenon remains unclear, but future research using accurate seizure and IES rate data streams in ambulatory subjects will enable further investigation into long-term temporal dynamics of IES and seizures and enable investigations exploring the IES rates^34^, changing IES morphology^41–43^ and circadian rhythms^35,44^ in association with seizures occurrence. The use of IES as a biomarker for seizure forecasting in the setting of EBS is an important direction for future investigation.

Regarding seizure reporting there are two important observations. Similar to previous studies, we found that patients^9,45^ and pet owners often do not create reliable seizure diaries when compared to gold-standard seizure catalogs created from automated seizure detection algorithms applied to continuous iEEG. This is not surprising given that seizures can be subtle, can go unnoticed by caregivers, and patients are often amnestic for their seizures. This result highlights the potential challenge of optimizing EBS and medical therapy if arguably the most critical measure of epilepsy therapy outcome, seizure rates, are inaccurate. This may play a role in the long time required for therapy optimization with current FDA approved devices. Furthermore, we determined that only 56.13 ± 30.77 % of ECoG captured seizures are reported by patients, thus many electrographic seizures would not be available for informing EBS therapy adjustments. Whether the unreported ECoG electrographic seizures reflect amnestic episodes or are truly subclinical seizures without clinical symptoms is unclear and raises an interesting future avenue of investigation where automated seizure detections could trigger an automated patient assessment^46^ to probe mood, cognition, memory, and motor impairments during and around seizures.

The current study has several limitations. The technology layers deployed in the system described here are associated with additional patient burden given that three rechargeable devices (Fig. 1.; implantable device, CTM, and tablet computer)^14^ must be periodically charged. This is the primary reason that not all data is captured during a long study. Given the fact that seizures can be relatively rare events the accumulation of adequate statistics remains a fundamental challenge for epilepsy research.

In summary, we present results from a powerful system integrating a new investigational neural sensing and stimulation device with local and distributed computing that should prove useful for future investigation and optimization of EBS in drug resistant epilepsy. This research identifies areas for future research including bi-directional interfaces to enable ECoG event triggered assessments^47^, continuous behavioral state tracking^48^ and seizure forecasting and adaptive EBS therapy. Future implantable systems with greater device computational power and data storage capacity will enable smart sampling paradigms to buffer data, run embedded algorithms, and trigger alarms for therapy change, behavioral queries, and data transfer that should enhance understanding of behavior and brain activity.

## Abbreviations

ANT: Anterior Nucleus Thalamus
AUPRC: Area Under Precision Recall Curve
AUROC: Area Under Receiver Operator Curve
CNN: Convolutional Neural Network
EBS: Electrical Brain Stimulation
ECoG: Electrocorticography
FFT: Fast Fourier Transform
iEEG: intracranial Electroencephalography
IES: Interictal Epileptiform Spikes
LSTM: Long-Short-Term-Memory
NV: NeuroVista
PRC: Precision Recall Curve
RC+S^™^: investigational Medtronic Summit RC+S^™^
ROC: Receiver Operator Curve
STFT: Short Time Fourier Transform
TLE: Temporal Lobe Epilepsy

## Acknowledgments

We thank the people with epilepsy for participating in this research. We appreciate the partnership with pet owners who made the canine research possible. We appreciate the technical and patient-focused support provided by Cindy Nelson and Karla Crockett. We thank Certicon a.s. for use of CyberPSG tool for visual review of EEG and Medtronic Plc for providing the investigational Medtronic Summit RC+S^™^ devices used in this research. We thank the research and clinical partners that provided the NeuroVista Inc. data used for developing the seizure detector. This research benefited from the community expertise and resources made available by the NIH Open Mind Consortium NIH U24-NS113637 (https://openmind-consortium.github.io/)

